# Mechanoreceptor-mediated circuit regulates cold tolerance in *Caenorhabditis elegans*

**DOI:** 10.1101/673863

**Authors:** Natsune Takagaki, Akane Ohta, Kohei Ohnishi, Yohei Minakuchi, Atsushi Toyoda, Yuichiro Fujiwara, Atsushi Kuhara

**Author notes:** Please address correspondences and requests to A.K. & A.O. (A.K.), (A.O.). Teaser: Thermosensitive mechanoreceptors regulate cold tolerance in *Caenorhabditis elegans*.

## Abstract

*C. elegans* mechanoreceptors located in the ASG sensoryneuron have been found to sense temperature — a key trait for animal survival. Experimental loss of xanthine dehydrogenase (XDH-1) function in the AIN and AVJ interneurons resulted in reduced cold tolerance and atypical neuronal response to changes in temperature. These interneurons are synapse with upstream neurons such as the mechanoreceptor-expressing ASG. Ca^2+^ imaging revealed that ASG responsiveness to temperature change via mechanoreceptor DEG-1, a Degenerin/Epithelial Sodium Channel (DEG/ENaC), affects downstream AIN and AVJ circuits. Ectopic expression of DEG-1 in the ASE gustatory neuron resulted in acquisition of thermosensitivity, while electrophysiological analysis revealed that DEG-1 was involved in temperature sensation. Together, these results suggest that cold tolerance is regulated by mechanoreceptor-mediated circuit calculation.

## Introduction

Animal temperature detection has previously been studied with a focus on transient receptor potential (TRP) channels. TRPV1, for example, is known to detect regions of high temperature, while TRPA1 detects regions of low temperature (Dhaka, Viswanath et al., 2006). Regarding TRP-independent temperature detection pathways, G protein– coupled receptor (GPCR)/rhodopsin in drosophila may act as a temperature receptor able to modulate decision-making behavior (Shen, Kwon et al., 2011). Furthermore, receptor-type guanylyl cyclases (rGCs) in the nematode worm *C. elegans* are thought to function as temperature receptors in the AFD temperature-sensing neuron given that ectopic expression of rGCs can confer temperature-dependent responses to heterologous cells (Takeishi, Yu et al., 2016).

*C. elegans* is an ideal model for the study of neural circuitry underlying cold tolerance given its simple nervous system composed of only 302 neurons as well as the number of well-studied molecular and genetic approaches currently available (Brenner, 1974). Mutant experimentation has also been well-documented, allowing for the identification of key genes and determination of specific neuron action sites (Barr, 2003). Finally, *C. elegans* temperature response has been analyzed with respect to many phenomena, including dauer larva formation (Barr, 2003), thermotactic behavior (Ohta & Kuhara, 2013), and cold tolerance(Ohta, Ujisawa et al., 2014, Okahata, Ohta et al., 2016, Sonoda, Ohta et al., 2016, Ujisawa, Ohta et al., 2018).

Together, the literature suggests that *C. elegans* possesses an adaptive mechanism to tolerate cold external environments. Wild-type worms grown at 15°C can survive at a temperature of 2°C, whereas 20°C cultivated worms cannot (Figure 1A) (Ohta et al., 2014, Sonoda et al., 2016, Ujisawa et al., 2018). Nematode cold tolerance is a process that involves a number of tissues, including the ASJ and ADL sensory neurons, intestinal cells, sperm, and muscle cells (Ohta et al., 2014, Sonoda et al., 2016, Ujisawa et al., 2018). In terms of sequence and site, the process begins when temperature is detected by the ASJ and ADL neurons located in the head (Ohta et al., 2014, Ujisawa et al., 2018). Next, insulin is released from the ASJ and binds to insulin receptors in the intestine and nervous tissue (Ujisawa et al., 2018), which initiates steroid-hormonal signaling to the sperm. Sperm in turn modulate ASJ neuronal activity in a feedback-like manner (Ohta et al., 2014). Genes are later expressed that ultimately modify bodily lipid composition (Murray, Hayward et al., 2007), which is considered central to cold tolerance. However, this mechanism describes negative regulation while overlooking yet-unexplored positive regulation.

**Figure 1.**
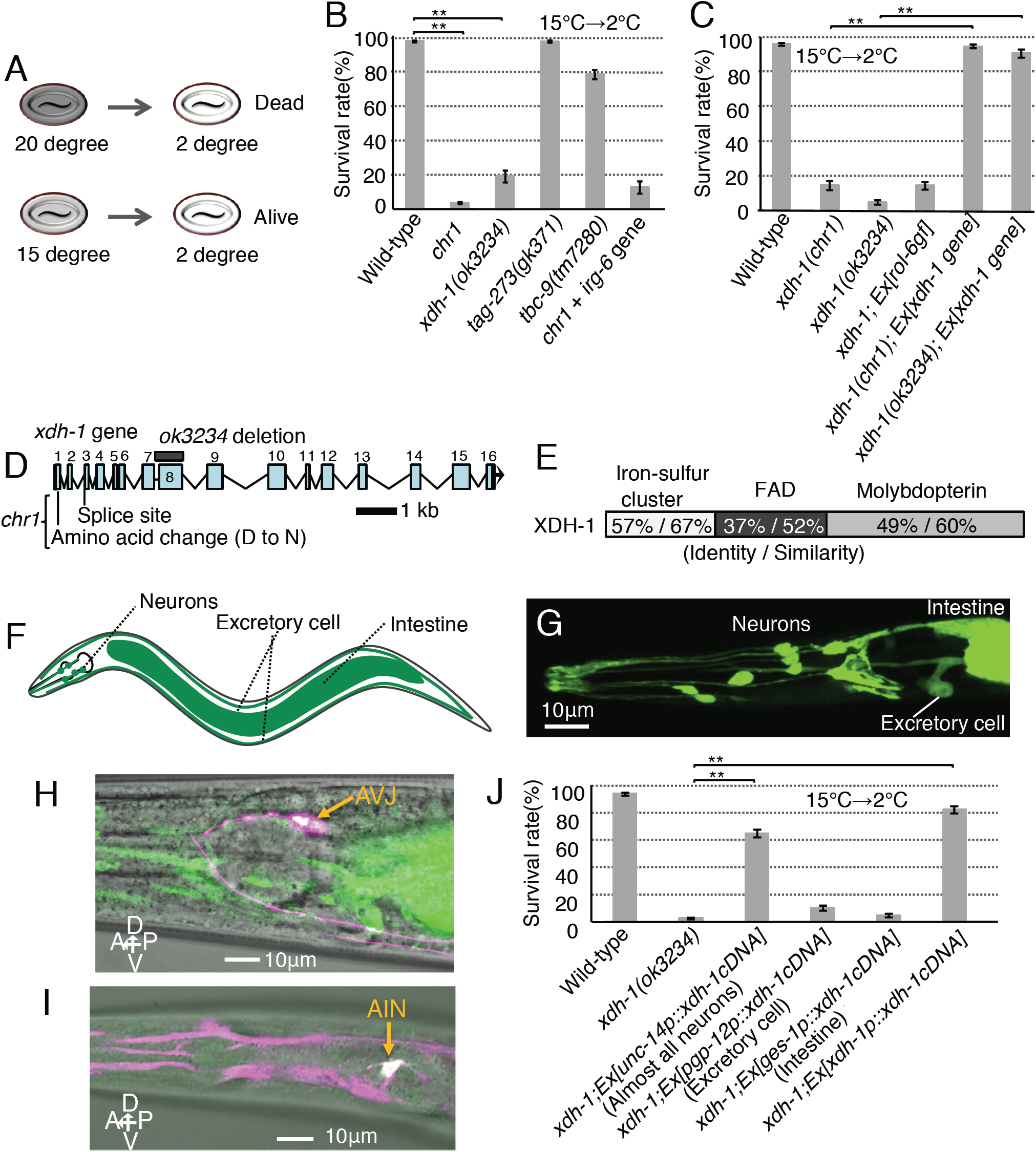
Neuronal XDH-1 regulates cold tolerance. Animals grown at 15°C were transferred to 2°C for 48 hours. Error bar indicates SEM (**B, C, J**). (**A**) Schematic of *C. elegans* cold tolerance. 20°C-cultivated worms do not survive at 2°C, while 15°C-cultivated worms do. (**B**) Cold tolerance when cultivated at 15°C. *chr1* and *xdh-1* mutants exhibit abnormal cold tolerance. Number of assays ≥ 10. Comparisons were performed using Dunnett’s test. **p < 0.01. (**C**) Transgenic rescue of *xdh-1* mutants with plasmid pNTN020 containing wild-type *xdh-1* fused with GFP and native promoter sequence. Number of assays ≥ 11. Comparisons were performed using the Tukey-Kramer method. **p < 0.01. (**D**) Exons of *xdh-1* gene are boxed and numbered. Positions of *chr1* point mutations and *ok3234* deletion mutation are shown. (**E**) *xdh-1* encodes xanthine dehydrogenase. Shading represents different domains in XDH-1. The amino acid identity and similarity between XDH-1 and human XDH are given for each domain. (**F**) Schematic diagram of expression pattern (green). (**G**) *xdh-1p::GFP* expressions in neurons, intestine, and excretory cells. (**H**) Wild-type animals expressing *xdh-1cDNA::GFP* driven by *xdh-1* promoter and AVJ-specific expression of dsRedm (wild-type; *Ex[xdh-1p::xdh-1cDNA::GFP, hlh-34p::dsRedm]*). Arrow indicates white region that suggests co-expression of GFP and dsRedm in AVJ interneuron. XDH-1::GFP fluorescence was spotty, and white region was faint. (**I**) Wild-type animals expressing dsRedm driven by *xdh-1* promoter and AIN-specific expression of YFP (wild-type; *Ex[xdh-1p::dsRedm, inx-17p::YFP]*). Arrow indicates white region that suggests co-expression of YFP and dsRedm in AIN interneuron. (**J**) Abnormal cold tolerance in *xdh-1* mutant was rescued by expressing *xdh-1cDNA* in almost all neurons. Number of assays ≥ 9. Comparisons were performed using the Tukey-Kramer method. **p < 0.01. Data from wild-type and xdh-1; *Ex[unc-14p::xdh-1cDNA]* animals are shown in Fig. 2A.

The <u>deg</u>enerin/<u>e</u>pithelial <u>Na</u>^+^ <u>c</u>hannel (DEG/ENaC) proteins comprise a diverse family of sodium ion channels (Chen, Bharill et al., 2016, Geffeney, Cueva et al., 2011, Waldmann, Champigny et al., 1996) involved in cellular functions such as mechanosensation (Geffeney et al., 2011, Zhong, Hwang et al., 2010), sour/salt taste (Chandrashekar, Kuhn et al., 2010, Liu, Leonard et al., 2003, Ugawa, Yamamoto et al., 2003), learning, memory, and synaptic plasticity (Wemmie, Askwith et al., 2003, Wemmie, Chen et al., 2002, Wemmie, Coryell et al., 2004, Zha, Wemmie et al., 2006, Ziemann, Allen et al., 2009). In mammals, the DEG/ENaC channel MDEG is abundantly expressed in the brain. (Waldmann et al., 1996), while the *C. elegans* homolog DEG-1 is expressed in association with multiple sensory neuron mechanoreceptors (Geffeney et al., 2011, Hall, Gu et al., 1997, Wang, Apicella et al., 2008). Although a decrease in temperature results in a change to the Na^+^ potential across MDEG (Askwith, Benson et al., 2001), it remains unknown whether DEG/ENaCs are directly involved in the temperature sensation process.

In mammals, xanthine oxidoreductase (XOR) is present as two interconvertible forms: xanthine dehydrogenase (XDH) and xanthine oxidase (XO). Moreover, both enzymes are active in the purine base salvage pathway. Notably, XDH converts hypoxanthine to xanthine while also oxidizing xanthine to urate. NADH is also produced simultaneous during the final step of purine salvage (Xi, Schneider et al., 2000).

XDH is expressed in the liver and small intestine (Chung, Baek et al., 1997). In humans, XDH expression levels are associated with tumor growth. Elevated expression of XDH is associated with tumor infiltration as well as upregulated proinflammatory and immune cytokine expression (Saidak, Louandre et al., 2018). XDH served as a useful biological parameter in one pan-cancer study (Saidak et al., 2018), yet the molecular function of XDH remains largely unexplored given that it may be either reversibly or irreversibly converted to XO in mammals (Saksela, Lapatto et al., 1999, Terawaki, Murase et al., 2017). Since invertebrate XOR is present only as XDH (Terawaki et al., 2017), *C. elegans* is thought to be a useful model.

We found that *xdh-1* mutant xanthine dehydrogenase knock-out worms exhibit abnormal cold tolerance, and that normal function can be recovered by expressing *xdh-1cDNA* in both AIN and AVJ interneurons. *In vivo* Ca^2+^ imaging revealed that XDH-1 acts as a positive temperature signal regulator in AIN and as a negative regulator in AVJ, and that temperature sensation by ASG sensoryneuron via mechanoreceptor DEG-1 affects neural activity of both AIN and AVJ interneurons. Ectopic expression of DEG-1 in the ASE, a non-thermosensitive chemosensory neuron, resulted in acquisition of temperature sensation. In addition, two-electrode voltage-clamp recording of *Xenopus* oocytes expressing DEG-1 demonstrated thermoreceptor-like behavior. These results together suggest that DEG-1, a DEG/ENaC type mechanoreceptor, acts as a temperature receptor and is required for the neural circuit calculation of the positive regulation of cold tolerance.

## Results and Discussion

To identify novel genes involved in cold tolerance, we isolated and analyzed the *chr1* mutation as characterized by decreased cold tolerance following cultivation at 15°C (Figures 1B and S1A-H, see supplemental material). Deep DNA sequencer and SNP analysis allowed the *chr1* mutation to be mapped from 6.13 cM to 16.24 cM on chromosome IV, where four genes were found to exhibit major mutations (Figures S1D-F) (accession number DRA: DRA 002599). We then evaluated cold tolerance in mutants for these genes (Figure 1B), with only *xdh-1* mutants exhibiting markedly abnormal cold tolerance after cultivation at 15°C. Moreover, abnormal cold tolerance in both *chr1* and *xdh-1* mutants was rescued by the expression of wild-type *xdh-1* (Figures 1C and D). These results together suggest that *xdh-1* is the primary gene responsible for the observed abnormal cold tolerance phenotype.

*xdh-1* encodes the *C. elegans* homolog of human xanthine dehydrogenase (XDH) (47% homologous) (Figures S2A and B), and XDH itself contains iron-sulfur clusters, FAD, and molybdopterin domains (Figures S2A-C). FAD is an NAD binding site, molybdopterin is a redox center, and XDH xanthine dehydrogenase in dimer form catalyzes the hydroxylation of xanthine as well as its subsequent conversion to uric acid (Figure S2C). Iron-sulfur cluster domains are heavily conserved throughout the animal kingdom (Figures S2A and B), and our *xdh-1(chr1)* mutants possessed two point-mutations in this domain: one in a conserved splicing acceptor site and another in a non-conserved amino acid residue (Figures 1D and S2B). Yet another allele, *ok3234*, contains a deletion mutation at an NAD binding site (Figures 1D and S2B), and we used this allele to conduct the following analysis.

Cells expressing XDH-1 located in the AVJ and AIN head nerves, the intestine, and excretory cells were analyzed by fluorescent protein expression driven by the *xdh-1* promoter (Figures 1F-I and S2D-F). To identify the essential tissue(s) responsible for *xdh-1*-dependent cold tolerance, we then expressed *xdh-1* cDNA in specific tissues (Figure 1J) and found that XDH-1 expression in nearly all neurons restored abnormal cold tolerance (Figure 1J; *xdh-1;Ex[unc-14p::xdh-1cDNA]*). *xdh-1cDNA* expression in intestine and excretory cells, however, did not rescue cold tolerance. This suggests neuronal XDH-1-activity is sufficient to maintain cold tolerance in *C. elegans*.

To determine the neuron type required for *xdh-1*-dependent cold tolerance, we performed a series of cell-specific rescue experiments by expressing *xdh-1cDNA* driven by various promoters (Figure 2). We found that abnormal *xdh-1* mutant cold tolerance was rescued by expressing *xdh-1cDNA* in multiple neurons, including the AIN and AVJ interneurons (Figure 2). Moreover, expression of *xdh-1cDNA* in AIN and AVJ interneurons simultaneously using *inx-17* and *hlh-34* promoters rescued cold tolerance in *xdh-1* mutants (Figure 2B). However, expression of XDH-1 in either AIN or AVJ alone did not rescue normal function (Figure 2B). These results suggest that XDH-1 expressed in both AIN and AVJ are required for cold tolerance.

**Figure 2.**
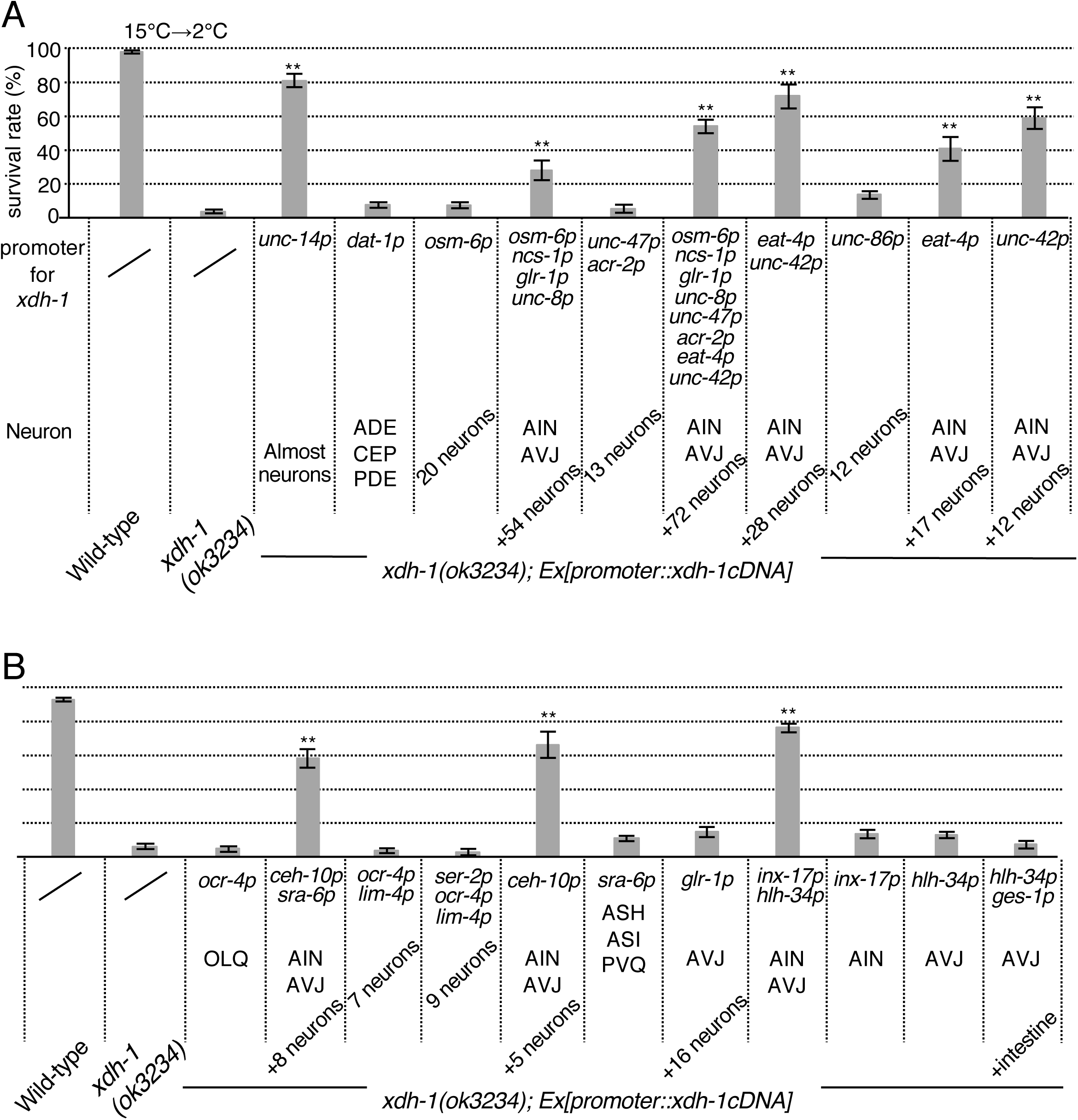
Cell-specific rescue experiments for abnormal xdh-1 mutant cold tolerance. (**A, B**) Expression of *xdh-1cDNA* driven by various promoters. 15°C-cultivated animals at adult stage were transferred to 2°C. Irregular cold tolerance in *xdh-1* mutants was rescued by expressing *xdh-1cDNA* in both AVJ and AIN interneurons. Number of assays ≥ 6. Error bar indicates SEM. Comparisons were performed using the Tukey-Kramer method. **p < 0.01. Data from wild-type and xdh-1; *Ex[unc-14p::xdh-1cDNA]* worms are shown in Fig. 1J.

Since AIN and AVJ are interneurons that receive a variety of sensory information, we hypothesized that temperature sensation by any upstream sensory neuron may affect the activity of AIN and/or AVJ. There are nine such neurons, and five are known to mechanoreceptor-expressing sensory neuron. To determine whether these mechanoreceptor neurons are involved in cold tolerance, we tested the cold tolerance of mutants defective in various aspects of mechano-transduction. Experimentation showed that mutation to any of a number of mechanoreceptor components could lead to abnormal cold tolerance (Figure 3A), but mutation to *deg-1*, which encodes a monomeric Degenerin/Epithelial Sodium Channel (DEG/ENaC)-type mechanoreceptor, resulted in particularly severe cold tolerance dysfunction (Figure 3A).

**Figure 3.**
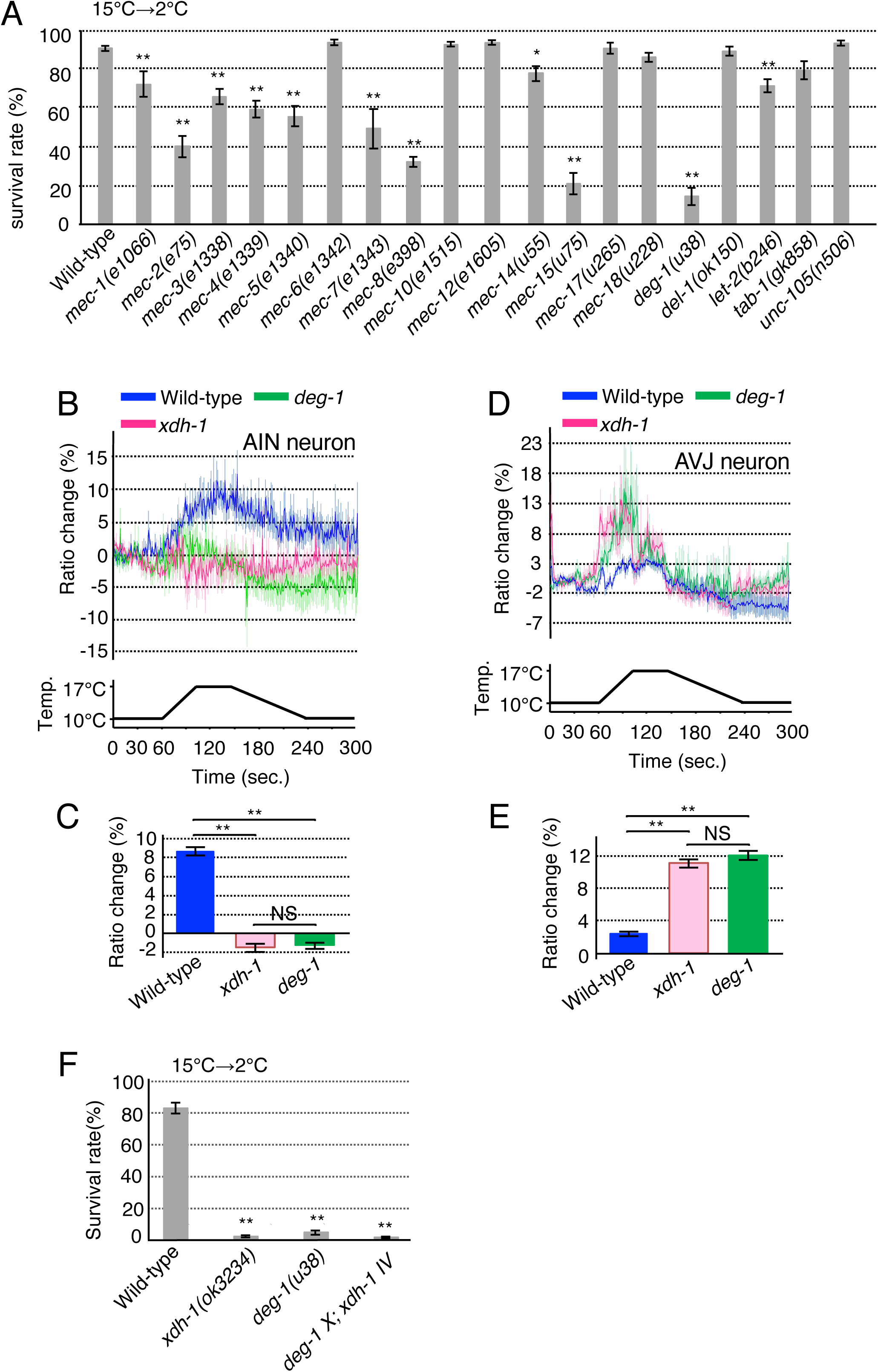
AIN and AVJ interneurons are involved in the thermosensitivity. Calcium imaging was performed using yellow cameleon 3.60 (**B to E**). Comparisons were performed using the Tukey-Kramer method. **p < 0.01. (**B, D, F**). (**A**) Cold tolerance of mechanoreceptor-related gene knock-out mutants. Animals grown at 15°C were transferred to 2°C for 96 hours. *deg-1* mutants exhibited pronounced abnormality. Number of assays ≥ 10. Error bar indicates SEM. Comparisons were performed using Dunnett’s test. *p < 0.05; **p < 0.01. (**B, C**) AIN calcium imaging in wild-type, *xdh-1* mutants and *deg-1* mutants cultivated at 15°C. Bar graph shows average change in cyan-yellow fluorescence ratio within 11 seconds from second 130 to 141. Each graph represents average response to temperature stimulus. Number of assays ≥ 17. Error bar indicates SEM. Bar graph color key is same as that of corresponding response curve in panel B (**C**). (**D, E**) AVJ calcium imaging in wild-type, *xdh-1* mutants and *deg-1* mutants cultivated at 15°C. Bar graph, average change in cyan-yellow fluorescence ratio within seconds 11 from second 90 to 101. Number of assays ≥ 17. Error bar indicates SEM. Bar graph color key is same as that of corresponding response curve in panel D (**E**). (**F**) Cold tolerance assays for *deg-1;xdh-1* double mutants. Number of assays ≥ 12. Error bar indicates SEM.

ASG is the sole sensory neuron pair upstream of AIN/AVJ that expresses DEG-1, and is located in the head of *C. elegans*. Ca^2+^ imaging revealed that wild-type ASG responds to temperature changes by increasing intracellular Ca^2+^ concentration (Figures 4A and B). In contrast, such a thermal response in *deg-1* mutants was lower than in wild-type, which was rescued by ASG-specific expression of *deg-1cDNA* (Figures 4A and B). These results suggest that DEG-1 is involved in the ASG sensory neuron temperature change response.

**Figure 4.**
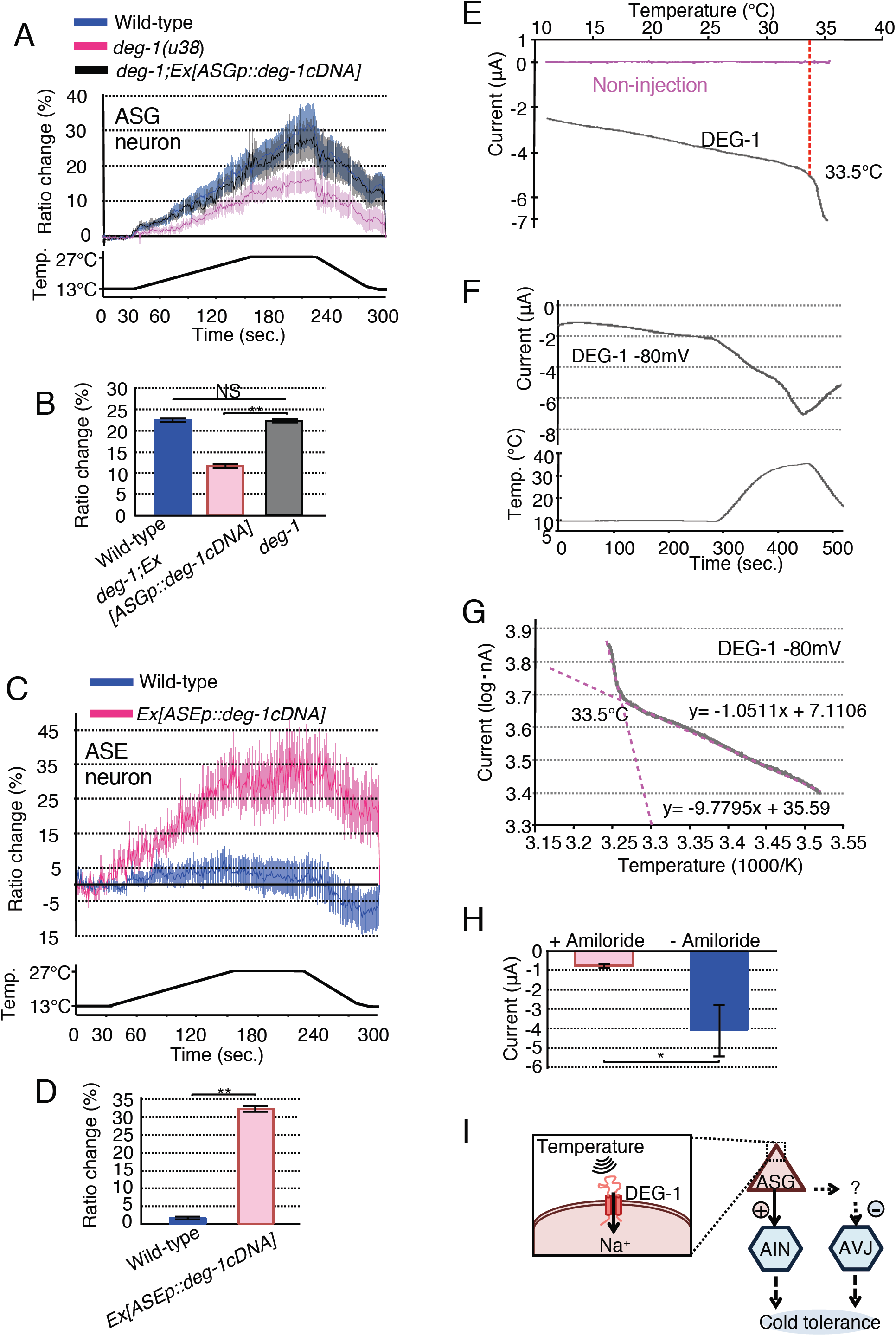
DEG-1 is involved in cold tolerance and thermosensitivity. Calcium imaging was performed using yellow cameleon 3.60 (**A, B**) and GCaMP8 with tag-RFP (**C, D**). Comparisons were performed using unpaired t test (Welch). *p < 0.05; **p < 0.01 (**D, H**). (**A, B**) Calcium imaging of ASG in animals cultivated at 15°C. Bar graph, average change in cyan-yellow fluorescence ratio within 11 seconds from second 230 to 241. Number of assays ≥ 20. Error bar indicates SEM. Previously reported, wild-type *deg-1* gene restored the *u38* abnormal touch sensitivity although the *u38* is a dominant negative mutation (Chalfie & Wolinsky, 1990) Bar graph color key is same as that of corresponding response curve in panel A (**B**). Comparisons were performed using the Tukey-Kramer method. **p < 0.01 (**B**). (**C, D**) Calcium imaging of non-thermosensitive ASE neurons in 15°C-cultivated wild-type worms ectopically expressing DEG-1 in ASE. Bar graph shows average change in cyan-yellow fluorescence ratio within 11 seconds from second 230 to 241. Number of assays ≥ 19. Error bar indicates SEM. Bar graph color key is same as that of corresponding response curve in panel C (**D**). (**E-H**) Reactions (representative current traces) to thermal stimulus in *Xenopus* oocytes expressing DEG-1. (**E**) Heating phase relationship between current and temperature shown in panels F. Data from non-injected oocytes (n = 8). (**F**) Representative current (upper) and temperature (lower) traces (n = 8). (**G**) Arrhenius plots from data in panels E. Data obtained from temperature increase is shown in panels F. Temperature threshold was determined by intersection of the two extended lines shown in magenta (n = 8). (**H**) Suppression of heat-evoked currents by incubating in bath’s solution with amiloride, an inhibitor of sodium ion channel (n ≥ 5). (**I**) A model for neuronal circuit for cold tolerance modulated by the signaling from ASG sensory neuron to AIN and AVJ interneuron. ASG senses temperature via DEG-1, connects to AIN (headed arrow). ASG indirectly connects to AVJ via unidentified neuron (dotted-line arrow). ASG-AIN-AVJ neural circuit positively regulates cold tolerance, in which ASG positively and negatively controls neuronal activity of AIN and AVJ respectively.

To investigate whether defective temperature sensation of ASG in *deg-1* mutants causes abnormal neuronal activity in its downstream interneurons AIN and AVJ, we performed calcium imaging using cameleon calcium indicator. Mutants *deg-1* AIN and AVJ calcium concentrations varied abnormally with thermal stimulus compared to wild-type (Figures 3B-E). In *deg-1* mutants, AIN activity diminished while AVJ activity increased (Figures 3B-E), suggesting that AIN and AVJ are both required for normal cold tolerance and that their actions oppose one another. Moreover, the responsiveness of *xdh-1* mutants AIN and AVJ were remarkably similar to the abnormal neural activities of these neurons in *deg-1* mutant, indicating that the neural circuit from ASG to AIN and AVJ neurons regulates cold tolerance. These results are consistent with a result of genetic epistasis between *xdh-1* and *deg-1* mutations that *deg-1;xdh-1* double mutants exhibited phenotypes comparable to either single mutation (Figure 3F), suggesting that *xdh-1* and *deg-1* act within the same pathway.

To determine whether mechanoreceptor DEG-1 is involved in temperature sensation, we ectopically expressed DEG-1 in the non-thermosensitive ASE gustatory neuron then measured the resulting intracellular calcium dynamics. ASE gustatory neurons ectopically expressing DEG-1 strongly responded to changes in temperature, while wild-type ASE did not (Figures 4C and D). These results suggest that ectopic expression of DEG-1 is sufficient to confer temperature responsiveness to the ASE gustatory neuron.

Next, we conducted electrophysiological experiments to determine the thermosensitivity of DEG-1 by applying the two-electrode voltage clamp recording method to *Xenopus* oocytes (Figures 4E-H). The DEG-1 mechanoreceptor and its mammalian homologue MDEG were expressed separately in *Xenopus* oocytes by injecting corresponding cRNAs for *deg-1cDNA* and *MDEGcDNA*. Thermal stimulus evoked internally-directed current in oocytes injected with *deg-1cRNA* and *MDEGcRNA* (Figures 4E-F, S4C and D), while control oocytes underwent no such changes (Figures 4E and S4C). These results suggest that DEG-1 and MDEG act as temperature-sensitive channels (Figures 4G and S4E; temperature threshold: 32.0 ± 0.8 °C for DEG-1 (N=8) and 31.0 ± 0.3 °C for MDEG (N=8)). It should be noted that the nematode vital temperature is between 13 and 27°C, while the *Xenopus* oocyte DEG-1 reactions took place at 32°C. Since the cold tolerant status in the worm body is constructed during the ambient cultivation condition such as 15 °C or 25 °C that is non-cold condition, as previously reported (Ohta et al., 2014), although we observed cold tolerance. Then, this is consistent with the result in this study that DEG-1 plays a role in temperature receptor sensing ambient temperature and in non-cold sensor.

Overall, the experiments performed in this study suggest that DEG/ENaC-type mechanoreceptor DEG-1 acts as a temperature sensor in ASG sensory neuron, which regulate AIN and AVJ interneurons to accomplish cold tolerance (Figure 4I).

## Materials and Methods

### Strains

We used the following *C. elegans* strains: N2 Bristol England (as wild-type) in all experiments, KHR066/RB2575 *flp-17(ok3587) / xdh-1(chr1)*, KHR067/RB2379 *xdh-1/F55B11.1(ok3234)*, VC883 *tag-273(gk371)*, FX07280 / *tbc-9(tm7280)*, KHR069 *xdh-1(chr1)*, CB1066 *mec-1(e1066)*, CB75 *mec-2(e75)*, CB1338 *mec-3(e1338)*, CB1339 *mec-4(e1339)*, CB1340 *mec-5(e1340)*, CB1472 *mec-6(e1342)*, CB2477 *mec-7(e1343)*, CB398 *mec-8(e398)*, CB1515 *mec-10(e1515)*, CB3284 *mec-12(e1605)*, TU55 *mec-14(u55)*, TU75 *mec-15(u75)*, TU265 *mec-17(u265)*, TU228 *mec-18(u228)*, TU38 *deg-1(u38)*, NC279 *del-1(ok150)*, DH246 *let-2(b246)*, VC1812 t*ab-1(gk858)*, and MT1098 *unc-105(n506)*. See supplemental material for further details.

### Cold-tolerance Assay

The cold-tolerance assay was performed according to previous reports (Ohta et al., 2014, Sonoda et al., 2016, Ujisawa et al., 2018, Ujisawa, Ohta et al., 2014). For this assay, we placed well-fed adult worms onto nematode growth medium (NGM) with 2% (w/v) agar, then seeded the medium with *Escherichia coli* OP50 once the worms began laying eggs. Adults were removed after 16-24 h at 15°C, and progeny were left to mature for 120-130 h at 15°C. Before the next generation had hatched, plates containing fresh adult worms were counted after being placed on ice for 20 minutes followed by transfer to a 2°C refrigerated cabinet (CRB-41A Hitachi, Japan) for 48-96 h. Temperature inside the refrigeration cabinet was monitored by both a digital and mercury thermometer. After the cold stimulus, plates were either repeatedly transferred to a room-temperature environment (total time over 3 hours) or stored at 15°C overnight. We then counted living and dead worms to calculate survival rates.

### Statistical analysis

Cold tolerance testing was conducted on 6 or more plates for 3 or more non-consecutive days. All error bars indicate standard error of the mean (SEM). All statistical analyses assume normal distribution and were performed using parametric tests, the Tukey-Kramer method, Dunnett’s test, or the unpaired t test (Welch). Multiple comparisons were performed using one-way ANOVA tested using the Tukey-Kramer method and Dunnett’s test. Dunnett’s test was performed to compare the left-most bar graph groups with other groups. Comparison between other group pairs was performed using the unpaired t test (Welch). *p < 0.05; **p < 0.01.

### Molecular biology

pNTN020 *xdh-1p::xdh-1 genomic gene::gfp* contains the *xdh-1* full-length gene and a 3,346 bp segment upstream of *xdh-1* amplified from the wild-type genome by PCR. GFP was inserted into the *xdh-1* full-length gene, excluding stop codon. pNTN026 *xdh-1p::gfp* contains the 3,346 bp upstream promoter sequence and the 3’-UTR of *xdh-1* amplified by PCR from pNTN020. GFP was then inserted by pPDF95.75. The *xdh-1p (1772 bp)::xdh-1cDNA::gfp* fragment contains the 1,772 bp upstream promoter sequence, the *xdh-1* gene, and *xdh-1cDNA* amplified by PCR from pNTN058 (supplemental methods), which was used for the transgenic expression experiment. pNTN118 *xdh-1p::dsRedm* contains the *xdh-1* promoter and dsRedm.

pNTN027 contains a Kozak sequence, the *xdh1*cDNA that was amplified by PCR from the cDNA library, and the 3’-UTR of the *unc-54* gene. The promoter sequences, *unc-14p* (1.4 kb), *pgp-12p* (3.5 kb), *ges-1p* (3.3 kb), *xdh-1p* (3.4 kb), *dat-1p* (0.7 kb), *osm-6p* (2 kb), *ncs-1p* (3.1 kb), *glr-1p* (5.4 kb), *unc-8p* (4.2 kb), *unc-47p* (0.3 kb), *acr-2p* (3.4 kb), *eat-4p* (6.4 kb), *unc-42p* (3 kb), *unc-86p* (3.6 kb), *ocr-4p* (4.8 kb), *ceh-10p* (3.5 kb), *sra-6p* (3.8 kb), *lim-4p* (3.6 kb), *ser-2p* (4.1 kb), *inx-17p* (1.2 kb), and *hlh-34p* (2.5 kb) were inserted upstream of pNTN027 *xdh-1cDNA,* to create respectively, pNTN034, 035, 036, 046, 047, 048, 049, 050, 051, 052, 053, 054, 055, 057, 059, 060, 061, 063, 064, 067, and 068 plasmids for cellular experimentation. pNTN075 *hlh-34p::yc3.60* contains the 2.5 kb *hlh-34p* gene and the *yc3.60* gene. pNTN106 *gcy-5p::deg-1cDNA* contains the *gcy-5* promoter received from Dr. Iino. pNTN116 contains the *inx-17* promoter, the *yc3.60* gene, and the 3’-UTR of the *let-858* gene. pNTN123 *gcy-21p::yc3.60* contains the 1,403 bp upstream promoter sequence of the *gcy-21* gene amplified by PCR from the wild-type genome, which was created by replacing the *hlh-34p* gene of pNTN075 with *gcy-21p*. Previous reports described the *gcy-21p::GFP* construct containing the 1^st^ and 2^nd^ exons and 1^st^ intron as inducing expression of GFP strongly in ASG and weakly in ADL. However, the *gcy-21p::GFP* construct excluding all exons and introns induces GFP expression in ASG only. We therefore used *gcy-21p* as an ASG-specific promoter. pNTN126 *gcy-21p::deg-1cDNA* contains the 1,403 bp upstream promoter sequence of the *gcy-21* and the *deg-1cDNA*. pMIU34 *flp-6p::CeG-CaMP8* contains a 2,680 bp upstream promoter sequence for the *flp-6* gene (*G-CaMP8*) and is codon-optimized for *C. elegans* (*CeG-CaMP8*). *deg-1cDNA* was inserted into a pGEMHE vector containing *Xenopus* beta-globin 5’ and 3’ UTR for electrophysiological recording (pNTN119).

### Two-electrode voltage clamp recording in *Xenopus* oocytes

*deg-1cRNA* and *MDEGcRNA* were separately injected into oocytes and incubated at 18°C for 3-6 days before electrophysiological recordings were made. We held membrane potential at −80mV and recorded macroscopic current using the two-electrode voltage-clamp technique with a bath clamp amplifier (OC-725C; Warner Instruments, USA) and pClamp software (Molecular Devices, USA) in bath solution containing 100 mM NaCl, 2 mM MgCl_2_, and 10 mM HEPES (pH 7.3). Temperature stimulation was regulated using a lab-made temperature controller with a range of 10 to 35°C and was monitored by both a thermistor prove adjacent to the oocytes and a thermometer (DIGITAL THERMOMETER PTC-401; UNIQUE MEDICAL, Japan). Arrhenius plot indicates the current amplitude induced by temperature changes on the y-axis (log scale) versus the inverse of temperature on the x-axis (1000/K). Temperature thresholds were determined by the intersection of the two liner regions (magenta lines), and all thresholds were then averaged. As a negative control experiment, we used amiloride, an inhibitor of sodium ion channel, and oocytes were incubated at 18°C in bath’s solution with 500μM amiloride (Sigma-Aldrich) for 48 hr.

### *In vivo* Calcium Imaging

*In vivo* calcium imaging was performed according to previous reports (Kuhara, Ohnishi et al., 2011, Ohta et al., 2014, Ujisawa et al., 2018). We used yellow cameleon (yc3.60) and GCaMP8 as genetically encodable calcium indicators. When using GCaMP8, we co-expressed tag-RFP (pKOB006 *gcy-5p::tag-RFP*) in order to measure the fluorescence ratio between GCaMP and tagRFP(Kobayashi, Nakano et al., 2016). Worms expressing the calcium indicator in one or more neurons were glued to a 2% (w/v) agar pad on glass, immersed in M9 buffer, and covered by a cover glass. Fluorescence of cyan (CFP) and yellow (YFP) by YC3.60, or green (GCaMP8) and red (tag-RFP), was simultaneously captured using a EM-CCD camera EVOLVE512 (photometrics, USA). Changes in intracellular calcium concentrations were measured as the yellow/cyan fluorescence ratio for YC3.60 or green/red fluorescence ratio for GCaMP8. See supplemental experimental material for more detail.

## Acknowledgments

We thank I. Mori, H. R. Horvitz, C. I. Bargmann, D. Yan, Y. Jin, K. Ashrafi, D. H. Hall, L. Bianchi, Y. Iino, T. Nakatani, T. Ii, J. Burkhead, J. M. Kaplan, T. G. Kusakabe and S. Mitani for sharing DNA constructs and strains; the National Bioresource Project (Japan) and the *Caenorhabditis* Genetic Center for strains; T. Miura and K. Kanai for supporting phenotypic experiments or maintenance of experimental systems; and members of the Kuhara Laboratory for comments and stimulating discussion. We thank Mr. Eric Odle for English editing and proofreading of the manuscript. We would like to thank the staff of Comparative Genomics Laboratory at NIG for supporting genome sequencing.

## Author contributions

N.T., A.O., K.O., Y.M., A.T., Y.F. and A.K. performed the experiments; N.T., A.O., A.T., Y.F. and A.K. designed the experiments, interpreted the results, and wrote the final report.

## Data and material availability

All data required to evaluate study conclusions can be found in either the main text or supplemental materials. Requests for further information should be addressed to A.O. or A.K. Information regarding data, figures, or other research findings may be addressed to the corresponding authors.

---

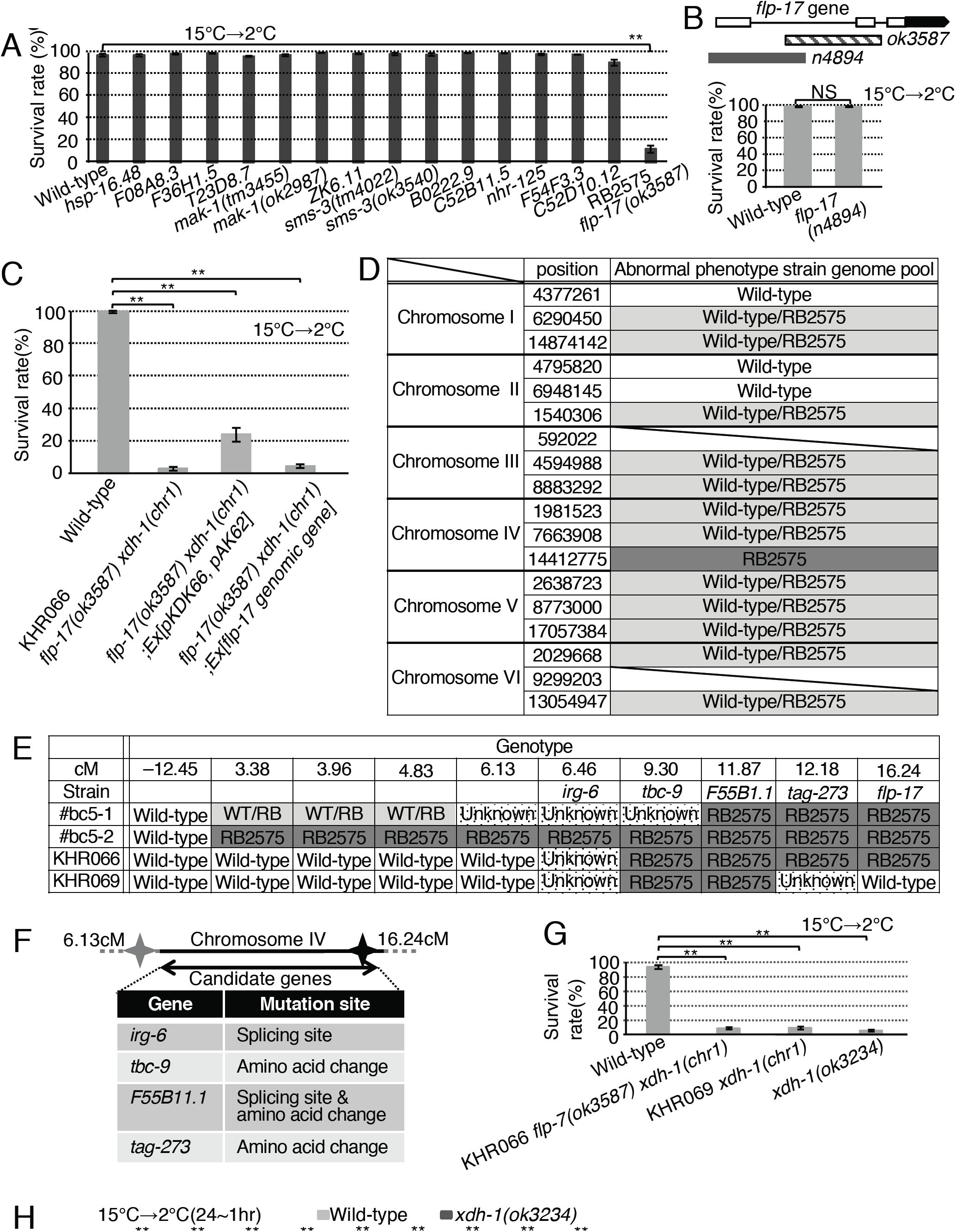

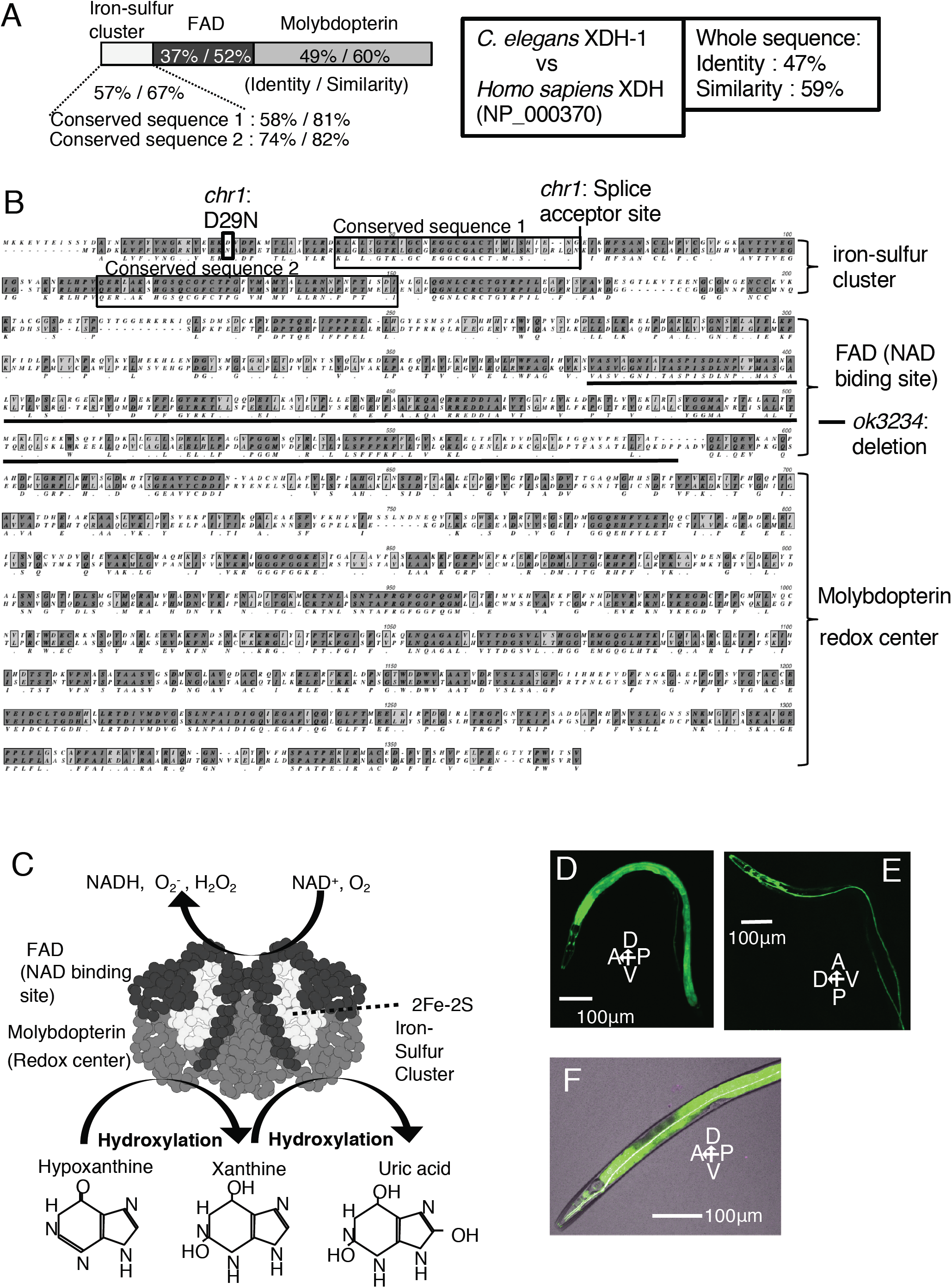

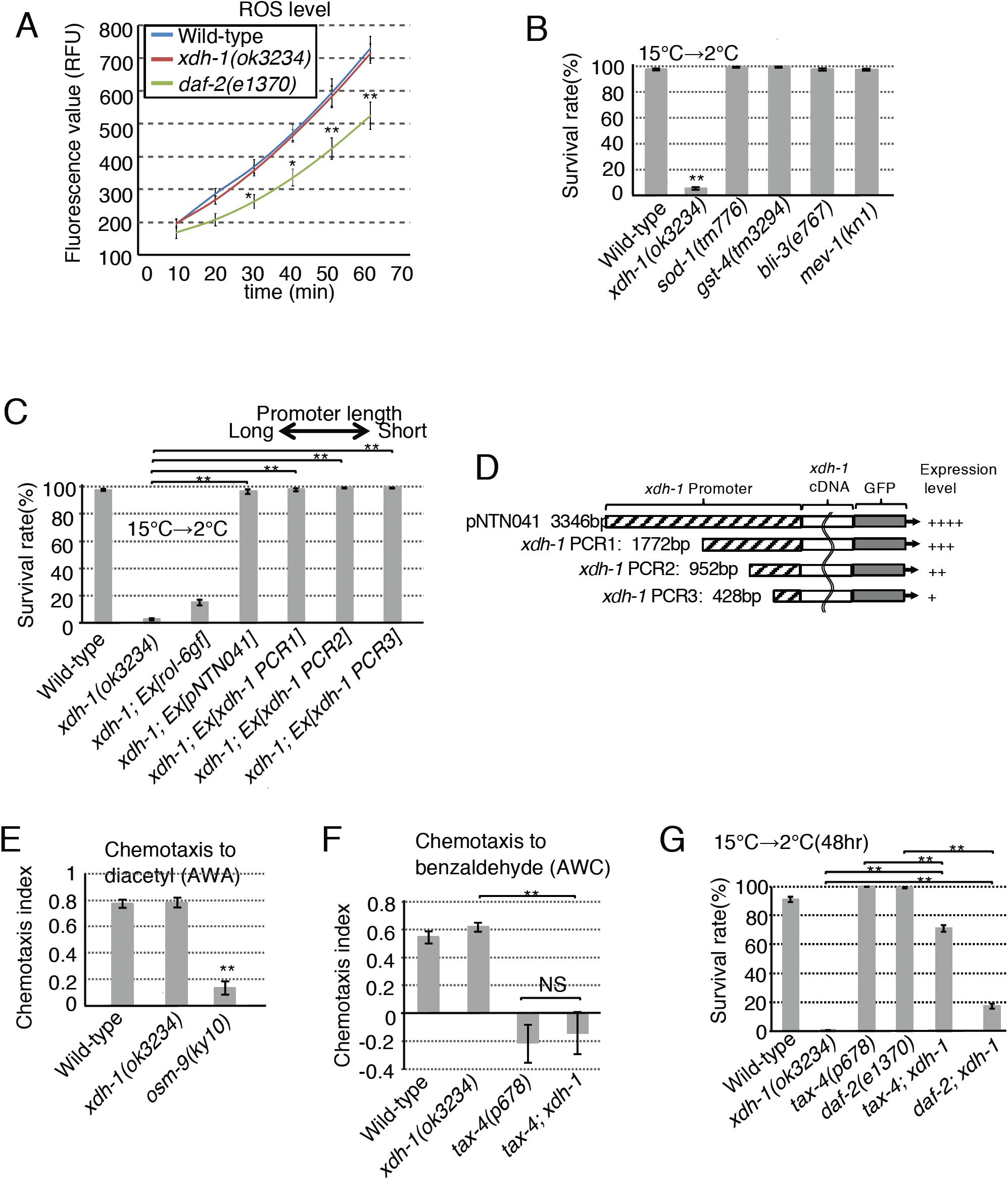

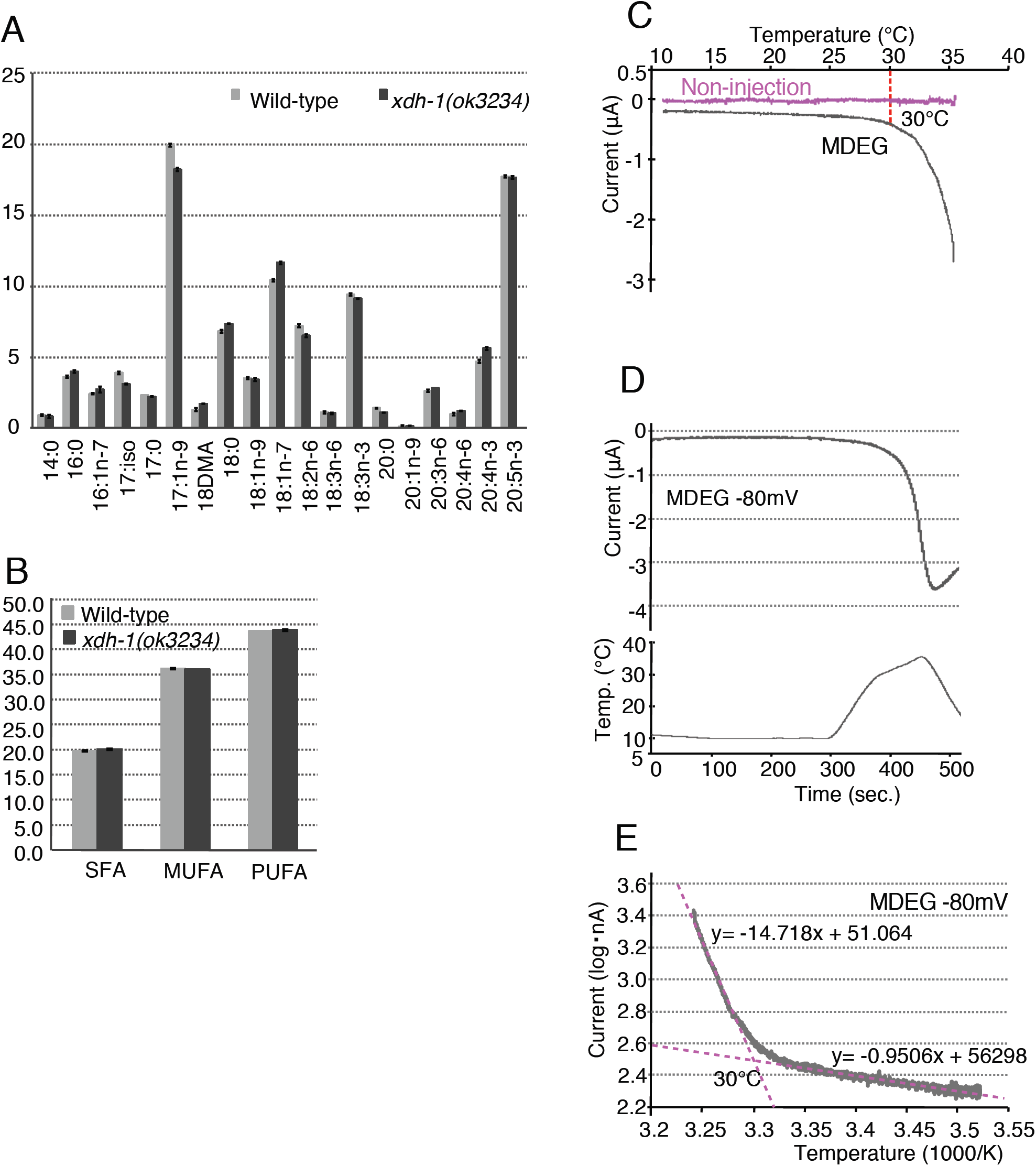

